# Tuning regimes in ant foraging dynamics depend on the existence of bistability

**DOI:** 10.1101/2025.07.22.666247

**Authors:** Colin M. Lynch, Bryan C. Daniels

## Abstract

Characterizing how behavior must be tuned to produce useful coordination is key to understanding the evolution and regulation of collective behavior. While computational models can answer this question for specific instances, recurring patterns in model dynamics hint at a more general means of classifying collective dynamics. Using ant foraging models as a foundational example, we investigate mechanisms that can produce symmetry-breaking transitions to bistability as a first basic classification of collective behavior. Collective transitions are functionally important: They lead to sudden changes in collective states, enhanced sensitivity to environmental inputs, and hysteresis. We use bifurcation theory to argue that the point at which discontinuous transitions merge at a continuous transition forms a codimension-2 bifurcation with universal properties, and that this point is functionally equivalent to the critical point of a phase diagram. We show how analogous bistable transitions appear across models of ant foraging with different mechanistic assumptions, and we explore how biologically relevant collective effects play out near the transition. This framework clarifies the difficulty of tuning collective behavior: locating a continuous transition typically requires tuning two parameters, while a discontinuous transition requires tuning only one. Finally, we explore conditions that degrade or destroy bistable transitions: heterogeneity blurs the transitions, while recruitment mechanisms that do not create a positive feedback loop do not display bistability at all.

## 1. Introduction

The macroscopic properties of biological systems emerge from the interactions of individual components, often without the guidance of a central controlling agent [1]. When the operations of individuals are performed jointly, the term *collective behavior* is used to describe the aggregate actions of these groups [2]. Collective behaviors occur across biological scales, at the level of genes [3, 4], neurons [5, 6], animal groups [7, 8], and human societies [9].

An important set of open questions in collective behavior concerns the evolution and tuning of decision-making algorithms used to coordinate group-level outcomes [10, 11, 12]. In cooperative systems, though individual components may have aligned interests, their ability to coordinate to produce useful behavior (for instance, interacting in a way that integrates information to improve collective decisions) depends on finding appropriate parameter regimes for both the behavior of individuals and the interactions between them. For example, house-hunting ants may adjust the rates at which they search for and accept candidate nests to trade off the speed of finding a new home versus the cost of committing to a suboptimal home [13]. This tuning of behavior is driven by evolutionary selection and active regulation that matches a group’s behavior to its environment [14]. The required specificity of tuning is key to understanding the difficulty of coordination, and likely depends on many details, including the complexity of the behavior and constraints on the mechanisms producing behavior [15]. Mathematically, we can describe this question as characterizing the mapping between tunable parameters and collective behavior, both in terms of the number of parameters needing to be tuned and the precision with which they must be tuned. An added complication is that many possible mappings may be able to produce similar behavior. In short, there are many ways that collective behavior could be successfully regulated, and we are searching for a categorization of these possible strategies that tells us something about their relative difficulty.

One promising characterization of collective strategies is in terms of collective transitions. Many living systems have been found to exist close to a transition at which one aggregate state becomes unstable [16, 17, 18]. Proximity to a transition can be adaptive [19], and biological systems are believed to regulate their position relative to such boundaries [20, 21, 22, 23]. Recent work has shown that heterogeneity among agents [24] and environmental feedback mechanisms [25] are key ingredients for tuning collective systems near these points.

An exciting possibility is that identifying collective transitions provides a general way to organize the space of possible collective behaviors and their difficulty of being tuned, one that generalizes across sets of distinct mechanistic details.^1^. Collective transitions can be studied using the concept of phase transitions in statistical physics [19, 29], and can also be approached by the related concept of bifurcations in dynamical systems theory [30, 31]. The concept of phase transitions was originally developed to describe phenomena such as the changing of a liquid into a crystalline solid at a particular temperature, for which the ordering of the crystal arises as the collective effect of interactions among enormous numbers of molecules. A key accomplishment of phase transition theory is to show how the unordered state becomes unstable to the development of order as a control parameter is varied. Bifurcations are a related concept that describes transitions between different attractors (dynamical regimes) in terms of changes to attractor stability as control parameters are varied. Transitions in living collectives occupy an awkward middle ground in which they do not fit comfortably within either of these frameworks. On the one hand, the theory of phase transitions most naturally handles systems that are in equilibrium and for which it is easy to scale up to extremely large numbers of components, neither of which is a very good match for biological systems. On the other hand, bifurcation theory is less convenient for incorporating stochasticity and does not particularly focus on many-body, collective effects. In this work, we aim to combine tools from both traditions in order to better clarify a ubiquitous form of collective transition — the transition to bistability — and to explore its biological implications.

The existence of well-developed classification schemes of bifurcations and phase transitions hints at the possibility that collective transitions might be classified in a similar way.^2^ This is promising because enumerating distinct transition types could offer (1) a way to make strong analogies between transitions happening across different collective systems; (2) a rationale for parsimonious phenomenological models that can be useful in systems that are difficult to model at an individual component scale [34]; (3) a better understanding of strategies for collective living — the typical patterns for how distributed living systems are organized and tuned in order to produce functional behavior [14].

As a first step in characterizing such transitions, the aggregate behavior of a collective can be summarized with some coarse-grained state variable (often termed an “order parameter” in statistical physics), e.g., an average over firing rates of individual neurons [35]. Some transitions result in a continuous change in this variable [36, 37], which is termed a continuous phase transition (sometimes called a second-order phase transition [38]). Examples include the change from free to congested flow in internet traffic [39], the effect of noise on collective cell migration [40], and environmental factors signaling a bee colony’s need to shift from worker to reproductive production [41]. Conversely, discontinuous phase transitions (or first-order transitions [38]) are abrupt and are associated with bistability and hysteresis [42], where system behavior depends on history [43]. Known cases include alarm-driven ant speed shifts [44, 45], foraging in Pharaoh ants [46], and shifts between ordered and disordered behavioral states in schools of fish [47].

Ants provide a uniquely tractable biological system for exploring the structure of collective transitions. Transitions can be reliably observed (such as the phases of nest migration, [48]), experimentally induced in laboratory settings (e.g., via alarm pheromones [49]), and monitored in the field at scale, including during phenomena like army ant bivouac formation [50]. Moreover, ants are ecologically dominant in many terrestrial ecosystems [51] and are presumed to be evolutionarily optimized for efficient resource allocation [52]. In harvester ants specifically, it has been well documented that foraging activity is regulated by the incoming rate of returning ants [53, 54, 55, 56]. This feedback system has been mathematically analyzed and shown to produce bifurcations and bistable dynamics in certain parameter regimes [57, 58]. However, this is only one of many mechanisms for foraging (reviewed in [59]). Other studies have implicated worker density in the nest as an important regulatory signal for activity modulation [60, 61]. Similarly, many existing models assume homogeneous individual behavior (i.e. [62]), despite findings that heterogeneity can significantly alter the dynamics of transitions [63]. While we focus on recruitment within ants, worker allocation strategies exist in other adaptive systems as well, including swarm robotics [64] and immune responses [65]. Therefore, our findings may not be unique to entomology.

These empirical insights from ant recruitment systems^3^ motivates a broader investigation into the dynamics of collective transitions, particularly how signal-mediated interactions give rise to emergent phenomena. Signal cascades lead to fundamentally collective responses: While solitary insects are individually able to perform tasks in an uncoordinated fashion, recruitment interactions create potentially useful coordination. Yet increasing interaction strength is not always better: for instance, amplified feedback signaling among immune cells can lead to a cytokine storm by triggering a self-reinforcing cascade of pro-inflammatory responses [71]. Intuition from statistical physics says that sufficiently strong interactions will lead to symmetry breaking into multiple possible collective states—in the simplest case, bistability. We ask here whether there are typical patterns in the way that bistability emerges in recruitment systems, how bistability can be tuned by regulatory or evolutionary dynamics, and the dynamical phenomena near the transition to bistability that are most functionally relevant.

We introduce three variants of a Markov-chain-based model of ant foraging to assess how different biological mechanisms contribute to the structure of collective transitions. The first is a flow-based model inspired by empirical results in harvester ants, where the return rate of ants triggers further activation [72]. In this model, the probability that an ant will leave the nest to forage depends on the rate of return of ants that have already foraged, as high rates are an indication that resources may be plentiful in the field. This model implicitly introduces positive feedback, as the more ants leave the nest, the more ants will return, increasing the probability that yet more workers will leave to forage.

The second is a density model, which investigates whether the spatial concentration of potential foragers within the nest alone can drive collective transitions. Here, the signal to forage depends on how many ants are currently in the nest. If there are many workers within the nest, then that may be a sign that there are not enough foragers, so the probability of leaving the nest increases. This model is implicitly dependent on negative feedback, as the more ants leave the nest, the less likely ants that are still in the nest will also leave. The third model is a modification of the first, in which individuals vary in their volatility (i.e. response thresholds) to the rate of returning ants. We examine the effects of heterogeneity under two conditions: one characterized by low variability and another by high variability. To test the universality of our results, we additionally analyze two supplementary formulations—an abstract Ising-like mean-field model and a more detailed existing stochastic dynamic model of this recruitment mechanism [25]. Both of these are also based on the returning-ant interaction mechanism, but span levels of abstraction to assess the robustness of the collective transition framework (details given in Supplemental information S1 and S2).

We analyze these models in terms of the dynamical properties of the group’s foraging rate and how they depend on tunable parameters. These dynamical properties, chosen to highlight aspects that are likely important to biological function, include sensitivity of the group’s foraging rate to changes in parameters (using a Fisher information measure), variability in foraging rate, and hysteresis. Together, these analyses allow us to examine how feedback and stochasticity determine the structure of collective behavior as a function of tunable parameters.

## 2. Methods: Simplified Foraging Models Based on Sigmoid Functions

Here, we define a simple set of models that represent the dynamics of foraging in the ant *Pogonomyrmex barbatus*. This framework is largely inspired by [25], but it has been simplified to expedite various measurements of collective dynamics. In section 2.1, we outline a discrete-time modeling framework as the basis of three specific models: a model in which the probability of foraging is dependent on the rate of workers entering the nest and all workers are the same (identical flow-in model), one in which workers are variable (heterogeneous flow-in model), and one in which workers are identical but the probability of leaving the nest depends on the number of workers within the nest (density model). Differences between these models are discussed in section 2.2.

See “Methods details” (section 5) for more details about computational and numerical methods.

### 2.1 Markovian Model Framework

In all three discrete time models, workers in a colony of size *N* are represented as a two state Markov chain, where workers are either residing in the nest (state 0) or they are actively foraging (state 1). *ρ*^0→1^ gives the probability of leaving the nest in a single timestep, and *ρ*^1→0^ is the probability that they will enter the nest^4^:

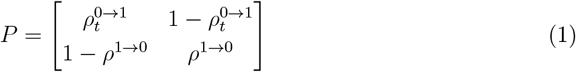

In timestep *t* ∈ {0, 1, … *τ*}, 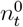 workers are foraging while 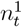 are inside the nest, thus 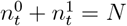. *ρ*^1→0^ is treated as a constant, whereas *ρ*^0→1^ depends on either the number of workers flowing into the nest at time 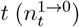 as is the case with the two flow-in models or it depends on the number of workers currently inside the nest 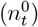 as is the case with the density model.

As there are only two possible states, the number of ants flowing into and out of the nest can be modeled as binomial random variables:

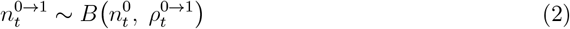

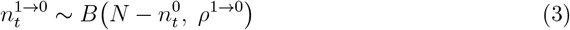

And since 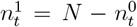, we can write the update function for the number of ants within the nest as:

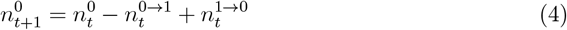

### 2.2 Probability of Leaving for Each Model

The three models differ only in the functional dependence of the probability of leaving the nest. In the flow-in models, foraging is stimulated by workers entering the nest [25], indicating that there is some external resource to the nest. These interactions set the probability 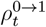, while *ρ*^1→0^ is determined by an individual’s ability to find food, which we assume is a constant so it does not depend on *t*.

*ρ*^0→1^ can be decomposed into two broad components: the propensity of the workers to forage even in the absence of any signal to do so, and the volatility of the available workers (that is, the willingness of workers to perform work given the inflow of other workers; Figure 1). The latter property can be further decomposed into different biologically-relevant features of workers, their sensitivity (their ability to resolve a signal), and the worker potential (the number of other workers in the nest needed to trigger a response). We assume that the probability of foraging increases monotonically with inflow as a sigmoid, using a modified version of the Fermi-Dirac distribution:

**Figure 1.**
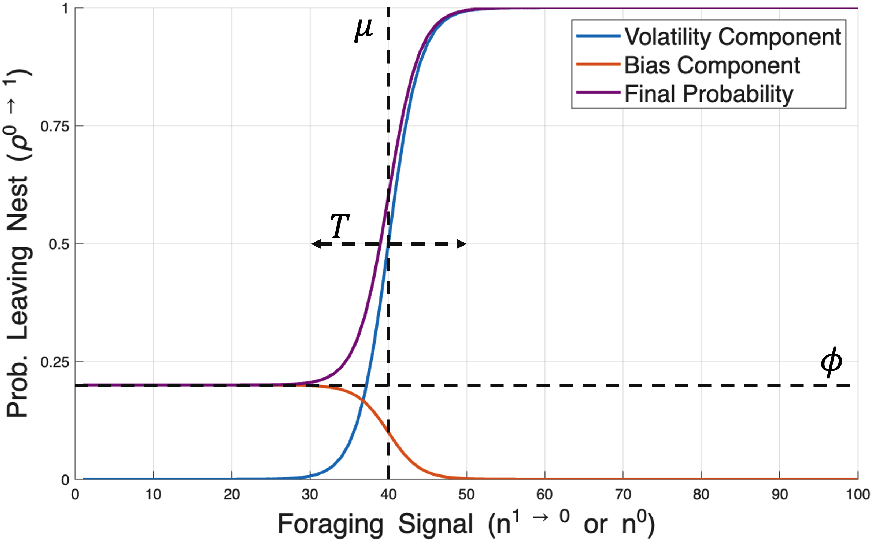
Social and non-social components contribute to the probability of foraging for flow-in models and the density model. The purple curve gives the probability of leaving the nest across different levels of the foraging signal. For flow-in models, this signal is the inflow rate. For the density model, this is the number of ants within the nest. This distribution is the sum of two other distributions: one representing the colony’s volatility in response to the work signal (blue line) and the other representing the propensity to forage in the absence of social signals (red line). The parameter *µ* gives the half-saturation point of the volatility component, *T* gives the width of the region between the minimum and maximum probability of leaving the nest, and *ϕ* gives the minimum probability.

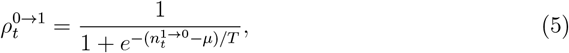

where *µ* ≥ 0 is the worker potential and *T >* 0 controls the sensitivity of the workers to the inflow cue. *µ* gives the half-saturation point of the probability curve (Figure 1), and lower values of *µ* make for a more responsive colony and are analogous to the total chemical potential. Low values of *T* result in a finely-resolved probability curve where workers are acutely aware of how many other workers are present in the nest. In contrast, high values of *T* reflect uncertainty in the worker, smoothing the probability curve, and thus operating like absolute temperature. The effects of the parameters on this equation are given in Supporting Information S3.

We assume a related form for the rate at which ants forage without social signals, which we refer to as foraging propensity:

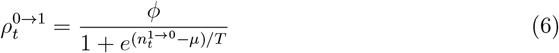

where *ϕ* is the individual foraging propensity. This function is at its maximum when equation 5 is at its minimum, so when the two are added together, it is setting the y-intercept of the sum while ensuring that the function outputs values between 0 and 1 (Figure 1):

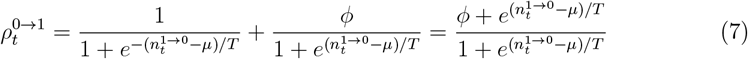

This equation produces probabilities when 0 ≤ *ϕ* ≤ 1.

In the density model, the only change to 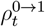 is to replace 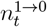 with 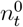:

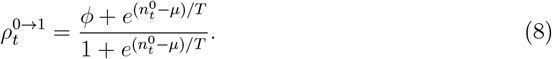

Finally, in the heterogeneous flow-in model, individuals vary in their foraging potential, which can also be understood as their threshold for foraging:

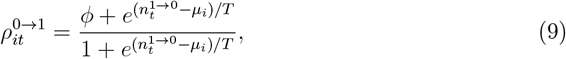

where

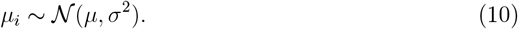

Units for each parameter are given in Table 1. Note that although the discrete-time structure of our model does not map directly onto real-time units, previous continuous-time models reported flow rates in units of ants per minute [25]. These models produced foraging rates in the range of 10–50 ants/min, comparable to those observed in our simulations (Figure 2). Based on this correspondence, we approximate each discrete timestep as representing one minute of real time.

**Table 1.** Description of variables used in each model. ^†^ High values of *ρ*^1→0^ in density model creates unrealistic oscillations within the model. We therefore show a case where *ρ*^1→0^ = 0.75 for a fair comparison to the identical flow-in model, but most simulations use *ρ*^1→0^ = 0.05. ^‡^*µ* also represents the mean of the normal distribution of potentials *µ*_*i*_ across individuals in the heterogeneous flow-in model.

| Variable | Variable Type | Range | Units | Parameter Selection |
| --- | --- | --- | --- | --- |
| Colony Size ($N$) | Constant | 100 | ants | NA |
| Number of Workers Outside of Nest ($n^1$) | State Variable | [0 $N$] | ants | NA |
| Number of Workers Inside of Nest ($n^0$) | State Variable | [0 $N$] | ants | NA |
| Initial Number of Workers Outside of Nest ($n_1^0$) | Free Parameter | [0 $N$] | ants | Drawn from uniform distribution |
| Number of Workers Entering Nest ($n^{0 \rightarrow 1}$) | State Variable | [0 $N$] | ants/min | NA |
| Number of Workers Leaving Nest ($n^{1 \rightarrow 0}$) | State Variable | [0 $N$] | ants/min | NA |
| Probability of Leaving Nest ($\rho^{0 \rightarrow 1}$) | State Variable | [0 1] | 1/min | NA |
| Probability of Entering Nest ($\rho^{1 \rightarrow 0}$) | Constant | $\in \{0.05, 0.75\}$ | 1/min | Either 0.75 for flow-In models, 0.05 or 0.75 for Density Model<sup>†</sup> |
| Total Number of Timesteps ($\tau$) | Constant | $\in \{100, 1000\}$ | Timesteps (≈ min) | Either 100 for most simulations or 1000 for tests of hysteresis |
| Worker Sensitivity to Foraging Signal ($T$) | Constant | 2 | flow-in: ants/min, density: ants | NA |
| Foraging Potential ($\mu$)<sup>‡</sup> | Free Parameter | [0 $N$] | 1/min | Systematically Iterated |
| Propensity to Forage in Absence of Signal ($\phi$) | Free Parameter | [0 1] | 1/min | Systematically Iterated |
| Amplitude of Foraging Potential ($\mathcal{A}$) | Constant | 20 | flow-in: ants/min, density: ants | NA |
| Standard Deviation in Foraging Potential ($\sigma$) | Free Parameter | $\in \{5, 20\}$ | 1/min | Used for two cases of heterogeneous flow-in models |
| Timestep ($t$) | Index | $\in \{1, 2, \dots, \tau\}$ | NA | NA |
| Individual ($i$) | Index | $\in \{1, 2, \dots, N\}$ | NA | NA |

**Figure 2.**
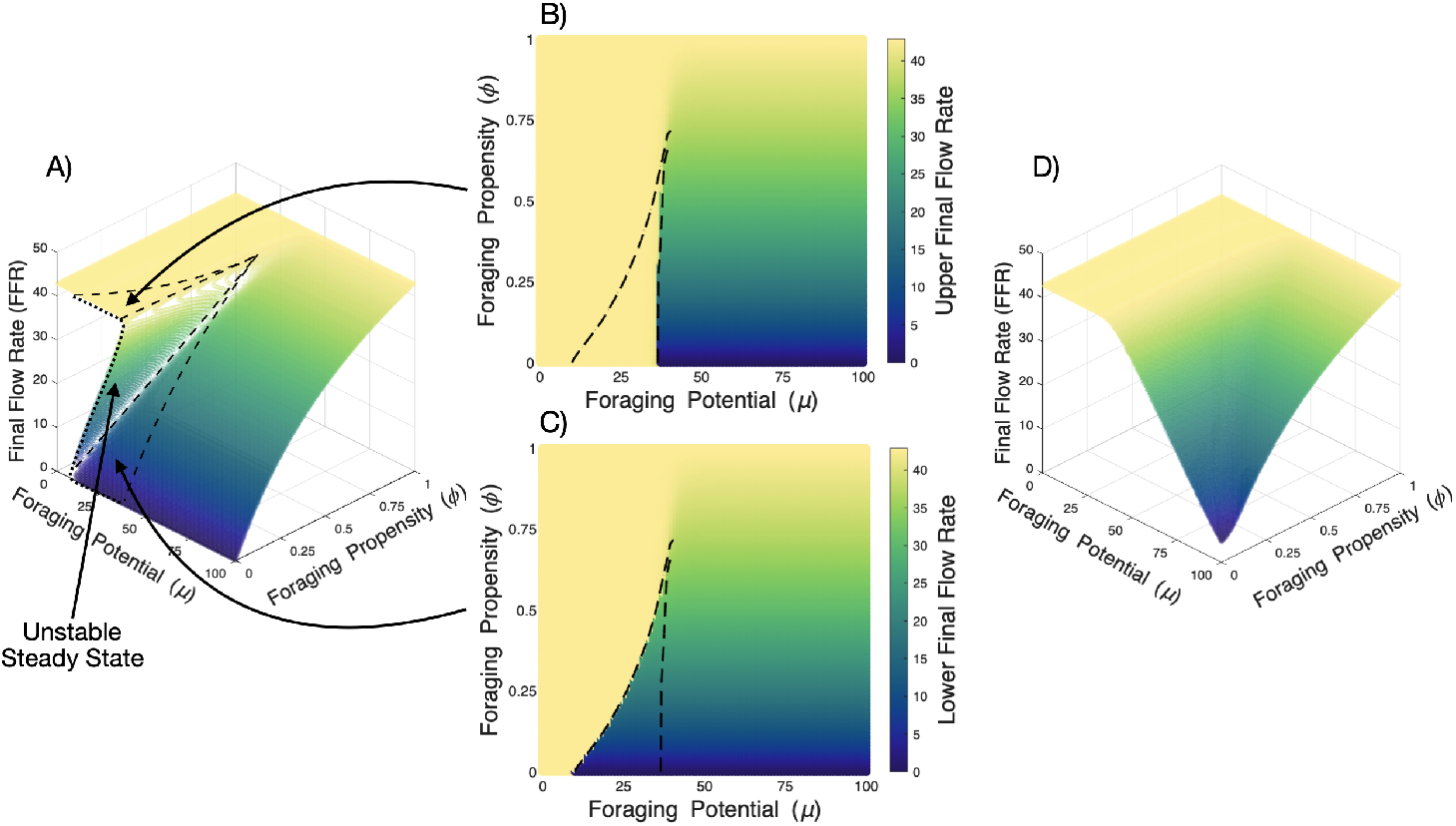
Bistability occurs in the flow-in model but not the density model. Final flow rates (steady state for the simulation) for the flow-in model across all combinations of the parameters *ϕ* and *µ* (A). In the region of the cusp bifurcation, the response surface is folded such that there are 3 possible steady states (the dashed lines show where this occurs). The middle values are unstable (Supporting Information S5), but the lower and upper flow rates are stable. They are therefore ignored when we identify the bistable region. To map the 3D plot onto a 2D plane, we can outline the bistable region where there is a stable upper steady state (B) and a stable lower steady state (C). Conversely, there is only a single steady state possible for the density model (D), so there is no bistable region. For all of these plots, *N* = 100, *T* = 2, and *ρ*^1→0^ = 0.75.

### 2.3 Measures of collective dynamics

To rigorously evaluate whether a model exhibits features consistent with phase transitions, it is essential to quantify multiple metrics of collective behavior. First, we measure the basin of attraction for different attractors. That is, in cases where there are multiple possible steady states, we calculate the probability that just one of these states will occur across a wide range of initial states. To quantify the variability of the collective rate of foraging, we compute two standard deviations: one that measures fluctuations in the rate across time within a given trial, and one that measures fluctuations in the rate across trials that start with different initial conditions. Hysteresis is an effect in which a transition happens at a different value as a parameter is increased as compared to when it is decreased. To quantify hysteresis, we systematically varied the response threshold *µ* for all individuals over time so that the second phase of the time series for *µ* is the mirror image of the first phase. We then measured foraging rates over that same time period, and measure the degree of asymmetry of foraging rates between the first and second phases. A high degree of asymmetry corresponds to high levels of hysteresis. To quantify the sensitivity of the collective rate of foraging *r* to various parameters controlling individual behavior, we compute multiple versions of the Fisher information. The Fisher information acts as a generalized measure of sensitivity [42], and it is known to diverge at continuous, equilibrium phase transitions [73]. In particular, the Fisher information measured with respect to a given parameter *a* measures how much the distribution of foraging rates *p*(*r*) changes as *a* is varied. Units for each collective outcome are given in Table 2.

**Table 2.** Outcome variables from models.

| Collective Outcome | Purpose | Description of Calculation | Units | Section |
| --- | --- | --- | --- | --- |
| Final Flow Rate | Rate of flow in and out of nest at steady state | Intersection of in- and out-flow rates | ants/min | 5.1 |
| Colony sensitivity to $T$ | Fisher information calculated wrt $T$ | Expected value of the observed information | flow-in model:<br>$\frac{\text{bits}}{(\text{ants/min})^2}$,<br>density:<br>bits/ants<sup>2</sup> | 5.2 |
| Colony sensitivity to $\rho^{0 \rightarrow 1}$ | Fisher information calculated wrt $\rho^{0 \rightarrow 1}$ | Expected value of the observed information | bits × min<sup>2</sup> | 5.2 |
| Variation Over Time | Variability of steady state for an initial set of conditions | Standard deviation of outflow rate over time | ants/min | 5.3 |
| Variation Over Initial Conditions | Sensitivity of model to initial conditions | Standard deviation of mean outflow rate across simulations | ants/min | 5.3 |
| Degree of Hysteresis | Asymmetry of foraging rates over time across the halfway point of simulation | 1 minus cross-correlation between first half and reversed second half of outflow rate over time | unitless | 5.4 |

## 3. Results

The cusp bifurcation, where two discontinuous transitions merge to form a continuous transition, is analogous to a critical point in statistical physics. To show how this 3-dimensional bifurcation diagram maps onto a 2-dimensional phase diagram, we first show all of the possible final flow rates for the identical flow-in model in Figure 2A. The point where the response surface starts folding over itself is the cusp, and this forms a region where there are 3 possible steady states, as opposed to all other regions, which are monostable. However, one steady point is unstable (Supporting Information S5), leaving only the top steady (Figure 2B) and the bottom steady (Figure 2C), creating a bistable region. The area outlined with dashed lines gives the bistable region, and the point where the two lines meet gives the critical point. In contrast, there is only one steady state possible in the density model (Figure 2D; Supporting Information S4), so there is no analogous critical point in this system.

### 3.1 Identical Flow-In Model

The cusp bifurcation drives collective dynamics in the phase space of colonies that use the flow-in mechanism for foraging. In the first column of Figure 3, we can see how the value of various collective dynamics changes across the phase space defined by the response threshold for foraging *µ* and the probability that an ant would forage in the absence of a foraging signal *ϕ*. The dashed lines show the bistable region (see Figure 2). In the first row, we estimate the sensitivity of the colony to small perturbations in the parameters, as measured by the Fisher information of the distribution of foraging rate with respect to *T* (main plot) and with respect to *ρ*^0→1^ (inlet plot). In both cases, the Fisher information is at its highest close to the discontinuous transitions and towards the continuous transition. At the edges of the bistable region, then, changes to parameters can result in discontinuous transitions in the foraging rate. In the second row, we estimate the degree of hysteresis of the colony, or how misaligned the second half of a simulation is with the first half given a symmetric change in the foraging potential (*µ*). We find that hysteresis is present only around the bistable region. In the third row, we measure the variability between simulations in foraging rates, which is highest in the middle of the bistable region. This marks the area of the phase space with the greatest sensitivity to initial conditions so that multiple states are possible when all other parameters are held constant. In the 4th row we measure the amount of variability in foraging rates within simulations, or temporal variability. It is highest near the edges of the cusp, as here colonies can hop between different states stochastically (Figure 5DEF). The rest of the parameter space consists of a monostable region, where parameter changes result in continuous changes to the foraging rate and nothing of interest occurs.

**Figure 3.**
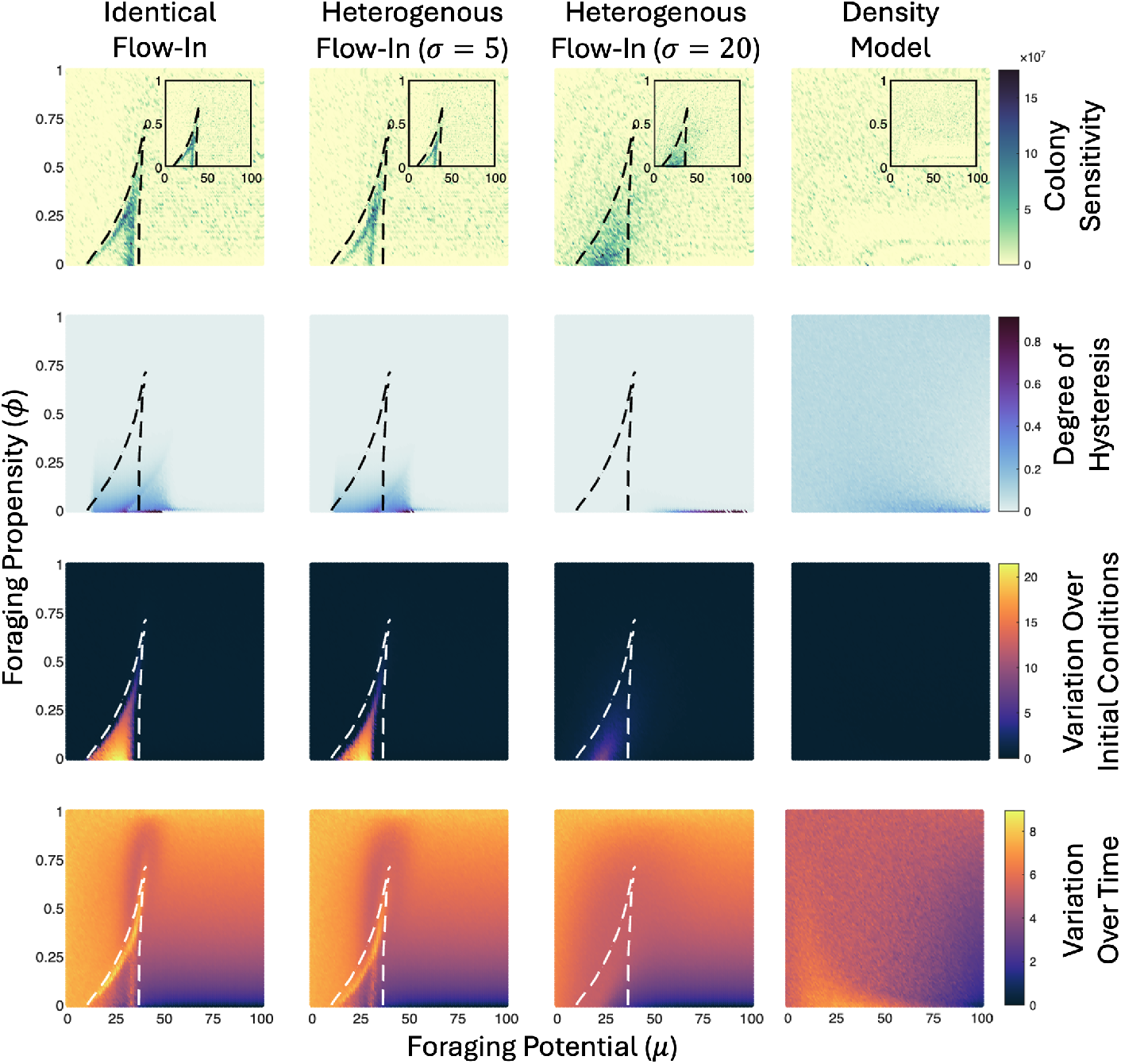
Bifurcations organize the phase space with respect to functionally relevant dynamical features in the flow-in models, but not the density model. Each panel displays how a collective outcome varies with the two parameters *µ* (horizontal axis) and *ϕ* (vertical axis). We measure four collective outcomes (rows) across four models (columns). Dashed lines indicate a bistable region as in Figure 2, with no bistable region for the density model. Colony sensitivity is measured as the Fisher information, calculated with respect to individual sensitivity *T* (main plots in the first row) and *ρ*^1→0^ (inset plots). The degree of hysteresis is measured as the asymmetry in foraging rates between the first and second half of simulations with time-varying *µ*. Variation over initial conditions is the standard deviation of the average flow rate across simulations, and variation over time is the average standard deviation of flow rates within simulations. Parameters are set as in Table 1.

### 3.2 Heterogeneous Flow-In Model

In the second and third columns of Figure 3, we present the phase diagrams for the heterogeneous flow-in models. Here, we present two versions of the heterogeneous flow-in model, one where the variability of the response threshold between individuals is low (second column, *σ* = 5) and one where it is high (third column, *σ* = 20). The phase diagrams from the low-variability colonies preserve many of the properties seen in the identical flow-in colonies (Figure 3, first column). However, many of these features tend to be canceled out by the high level of noise in the highly variable colonies. Still, many of these properties remain; they are just restricted to smaller, less well-defined domains.

### 3.3 Density Model

In the fourth column of Figure 3, we present the phase diagrams for the density model. In contrast to the flow-in models, changes to the system dynamics of the density model are more diffused across the parameter space. Fisher information (top row), calculated with respect to both *T* in the main plots and *ρ*^1→0^ in the inset plots, is lower in magnitude and less well defined for the density model compared to the flow-in models. The density model also always produces the same steady state, so variability across simulations is near 0 (third row). A similar pattern is observed for the degree of hysteresis (second row), which is likewise diminished and less distinct. Finally, temporal variability (fourth row) of the foraging rate varies across the phase space, but it does so more diffusely than in the flow-in model, lacking the sharp ridges in variability. Changing parameters for the density model, then, only results in changing the final flow rate (Figure 2D) and not much else.

### 3.4 Other models

Cusp-like bifurcations also occur in two additional flow-in-like ant foraging models discussed in Supporting Information: an Ising-like mean-field approximation and a more detailed model based on [25].

## 4. Discussion

To explore the space of possible collective dynamics and how they depend on tunable parameters, we analyzed multiple related models of ant foraging behavior. In one set of models, the signal to forage is the flow rate of workers entering the nest. In another model, the signal to forage is the density of workers within the nest. Despite their shared basis, these models differ markedly in how their collective dynamics depend on tunable parameters. In the flow rate models, positive feedback emerges from the fact that foraging rates themselves depend positively on foraging rates, so increasing the rate will result in uncommitted ants having a higher probability of foraging. The presence of this feedback results in bistability under some parameter regimes, where the foraging rate can quickly build to a high stable state or quickly decay to a lower stable state. The phase space for the flow rate model is further characterized by a cusp bifurcation that leads to dynamical effects including bistability [74], colony-level sensitivity to perturbations in parameters (also known as susceptibility in statistical physics; [75]), and hysteresis [76]. At the edges of the bistable region, small parameter changes can drive sharp shifts in the behavioral state of the colony. The generalized sensitivity measured by the Fisher information is large along these edges. In particular, the sensitivity is large near the continuous cusp bifurcation, as expected from criticality theory in statistical physics [73], and it is also large near the discontinuous saddle-node edges of the bistable region. Elsewhere in the phase diagram, only gradual changes are possible, so the flow rate models are highly tunable only near these transitions.

In contrast, colonies that use worker density as a signal for foraging cannot be tuned to the same degree. This mechanism lacks amplification and, indeed, exhibits negative feedback. Here, high density triggers further exploration, which then decreases the density, so that uncommitted ants are less likely to forage. This negative feedback promotes monostability and robustness to alterations in parameters [77]. The dynamics present in the flow rate models are not possible in the density model, and as a result, regions with distinct collective behavior are more dispersed in parameter space. Sensitivity and variability are present but weak. Tuning of the parameters, then, only results in small changes to the flow rate. The stubbornness that results from this negative feedback may make the colony unable to respond to important signals.

There are many potential functional benefits to social insect colonies by having dynamics near a bistability. First, as has been emphasized in the criticality literature, the maintenance of high Fisher information can be interpreted as enhancing the colony’s ability to react to environmental perturbations [78]. Second, there are many environmental contexts in which discrete, all-or-nothing responses could be adaptive, especially when intermediate levels of worker engagement are suboptimal. For example, a small number of individuals may be insufficient or inefficient for tasks such as collective transport [79] and the capturing of large prey [80]. In such cases, a bistable process that allocates either no workers or a large number could be preferable to one that allocates a highly variable number. Similarly, hysteresis is known to stabilize emergent collective structures such as self-assembled bridges [81].

Our characterization of collective transitions demonstrates how criticality theory might be usefully generalized to biological systems with large but not extremely large *N*: systems poised near both continuous and discontinuous transitions (bifurcations) are particularly sensitive, adaptive, and capable of transitioning between behavioral regimes with minimal input. These properties are thought to offer functional advantages in other biological contexts as well, including ecological communities [82] and abstract models of evolving behavior [83]. All-or-nothing responses are also thought to be useful in the sense of digital computing as implemented by, for example, neurons [84].

Still, these properties can only be adaptive if they can survive the noise inherent to any biological system [85]. Fortunately, we find that the flow rate models that incorporate variation among individuals retain many of the features of the identical model at low levels of individual variation. However, as heterogeneity increases, the sharpness of the collective transitions diminishes. This is consistent with the idea that disorder can smear transitions - similar to a Griffiths [86] - suggesting that there could be limits of tunability in the presence of noise. This finding aligns with work on disorder in complex systems and suggests that excessive heterogeneity may obscure the critical region [87, 86] ^5^.

Utility also depends on the ubiquity of these results. Do these properties emerge from arbitrary assumptions of the model, or do they emerge from deeper principles? Notably, we observed similar bifurcation structures in two additional models: an Ising-like mean-field approximation and a continuous Fitzhugh-Nagumo model derived from [25] (see Supporting Information). Despite their differences in complexity and formulation as well as the types of parameters that were varied, both models produce transitions to bistability that result from cusp-like bifurcations. In the Ising-like model, two saddle-node bifurcations merge to form the more familiar version of a cusp. In the model from [25], a saddle-node bifurcation appears to merge with a transcritical bifurcation to form a bistable region (see Figure S3). Beyond these, at least two further models of ant foraging in the literature, built using different mathematical structures, also produce bifurcations with bistable regions [57, 58, 90]. This invariance to the details of the mechanisms suggests that the presence of these bifurcations is not merely an artifact of potentially arbitrary modeling choices, but rather emerges from the potential for positive feedback. In this way, we can envision usefully classifying collective mechanisms in terms of the bifurcation structures they can create.

The construction of phase diagrams in each model proceeds from mean-field self-consistency relationships that govern steady-state behavior. In our case, the input/output functions can be adequately represented by (cubic) polynomials, even in systems lacking closed-form analytical solutions (see Supporting Information S2), allowing for trivial analysis of the system. A cubic form of the mean-field input/output function can create regions of positive feedback and thereby naturally leads to bistability and cusp bifurcations. Part of the reason the cubic polynomial is a good fit here is due to the finite population of a colony. The outgoing foraging rate cannot increase indefinitely, as the flow rate cannot exceed colony size, so eventually the outgoing rate must trend back downwards for large inflow rates. For some parameters, this initial increase, and then decrease of the output function allows for three crossings along the input/output identity line, allowing for bistability. However, it is also possible to generate a self-consistency curve with 3 crossings with an infinite population. An Ising-like model of this foraging mechanism has a self-consistency curve that takes the form of a hyperbolic tangent (Supporting Information S1).

The cubic shape of the self-consistency curve (which has been observed in other models, i.e. [46]) can be controlled by just two parameters: the threshold to social inputs and the probability of foraging without social inputs. Therefore, tuning of just these two parameters is sufficient to locate collective transitions across all the models. This low-dimensional structure is evident, too, in models of differing complexity: each Markovian model from the main text, as well as a simpler mean field model and a more complicated dynamical model explored in Supporting Information. At each level of complexity, we find analogous bifurcations in a two-dimensional parameter space. According to bifurcation theory, this is not unexpected [91]. In particular, the edges of the bistable region in our identical model correspond to saddle-node bifurcations, one of the simplest and most ubiquitous bifurcations in dynamical systems. A saddle-node bifurcation has codimension 1: that is, one parameter typically needs to be tuned to find the bifurcation. Starting from an existing saddle-node bifurcation and following it as a second parameter is varied, as along the sides of the bistable regions in our phase diagrams, can locate bifurcations of codimension 2. In our case, we find the simplest type of codimension 2 bifurcation: two saddle-node bifurcations join at a cusp. Given this picture, for a generic transition to bistability, we might expect that (1) without further tuning, the transition is likely to be discontinuous; (2) the ability to tune a second parameter is likely to be enough to find a set of parameters corresponding to a continuous transition. This is indeed the result that we find in all the ant foraging models that support positive feedback. Note that this argument works for any parameter that is available for tuning. If there are a number *N*_*p*_ of such parameters, we can envision system dynamics as being specified by a point in *N*_*p*_-dimensional parameter space, and tuning two parameters corresponds to taking a 2D slice out of this larger space. A bifurcation with codimension *c* will generally occur not along a single line in this larger space, but on a manifold of dimension *N*_*p*_ − *c*. For this reason, if a transition of codimension *c* exists, varying *c* independent parameters will be sufficient to locate such a bifurcation except in special cases of parameters that are orthogonal to this manifold. In this way, the codimension of a bifurcation effectively sets the “difficulty” of finding it—with relevance to the regulation of collective behavior on short timescales and its evolution on longer timescales.

This approach of characterizing bifurcations constitutes a theoretical framework that could be used as a basis for experimental predictions. For example, empirical phase diagrams could be constructed by systematically varying environmental parameters and measuring resulting foraging rates. Low humidity has been shown to suppress foraging activity in harvester ants by increasing nest bias [92], which could correspond to a uni-directional shift in parameter space. Similarly, dopamine levels affect foraging propensity and volatility [93], and may modulate parameters akin to volatility or threshold variance in our models. By altering these parameters in controlled experiments, our models predict collective transitions that may be visible in the dynamics of foraging rate.

Our results here are one step toward a more general classification scheme for collective transitions in living systems. Identifying the potential types of transitions and the mechanisms that produce them will be a crucial ingredient of understanding the evolution and tuning of collective behavior. In particular, categorizing mechanisms by the transitions they can support, characterizing the types of feedback they generate, and classifying transitions by codimension helps to predict the relative difficulty in finding and regulating functional collective behavior through the tuning of underlying parameters. We envision that this simplifying framework will be useful more broadly within studies of communication networks and informational cascades across many parts of biology and society, including social insects (including alarm and other signaling cascades), neuroscience (where there is already a vibrant literature analyzing such collective transitions), and perhaps even human information networks.

## 5. Methods details

In section 5.1, we show how we numerically approximate the steady states of the identical flow-in model and the density model. In section 5.2, we derive the Fisher information of each model with respect to two different parameters. In section 5.3, we describe how we run simulations so that we can measure across and within simulation variation of a state variable as well as the probability that different stable states will result for different parameter combinations. In section 5.4, we introduce a time dependency of one parameter to quantify hysteresis. See Table 1 for a description of each of the state variables, constants, and free parameters of each model, and Table 2 for a description of each collective output.

### 5.1 Numerical Approximation of Steady State

At a steady state fixed point, the inflow of ants matches the outflow. That is:

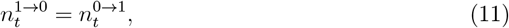

which on average is equal to

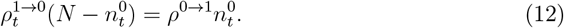

However, due to the presence of the exponential function in 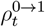 for both the flow-in and density models, this is a transcendental equation which cannot be solved analytically. Instead we use a numerical approach to find the final flow rate (steady state) for the identical flow-in model and the density model. This approach fails for the heterogeneous flow-in model, and so will not be attempted.

First, we find the outflow rate for the flow-in model and the density model, evaluated over all possible values of workers waiting within the nest 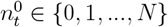:

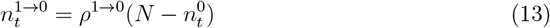

We then find the roots within a small tolerance limit:

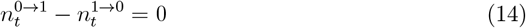

Each root is a final flow rate 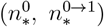.

As the outflow rate of the identical flow-in model can be cubic as a function of inflow (Figure 4; Supporting Information S2), there are up to 3 possible final flow rates. When there is only 1 final flow rate, the steady state is stable (closed circle). When there are 3 final flow rates, one of these points is unstable (open circle). While it is possible to only have 2 final flow rates in this system, this possibility is extremely rare and only results in a single stable steady state, occurring only at the cusp bifurcation. Additionally, the exact shape of the self-consistency curve influences the probability that different steady states will occur across random conditions (Figure 2).

**Figure 4.**
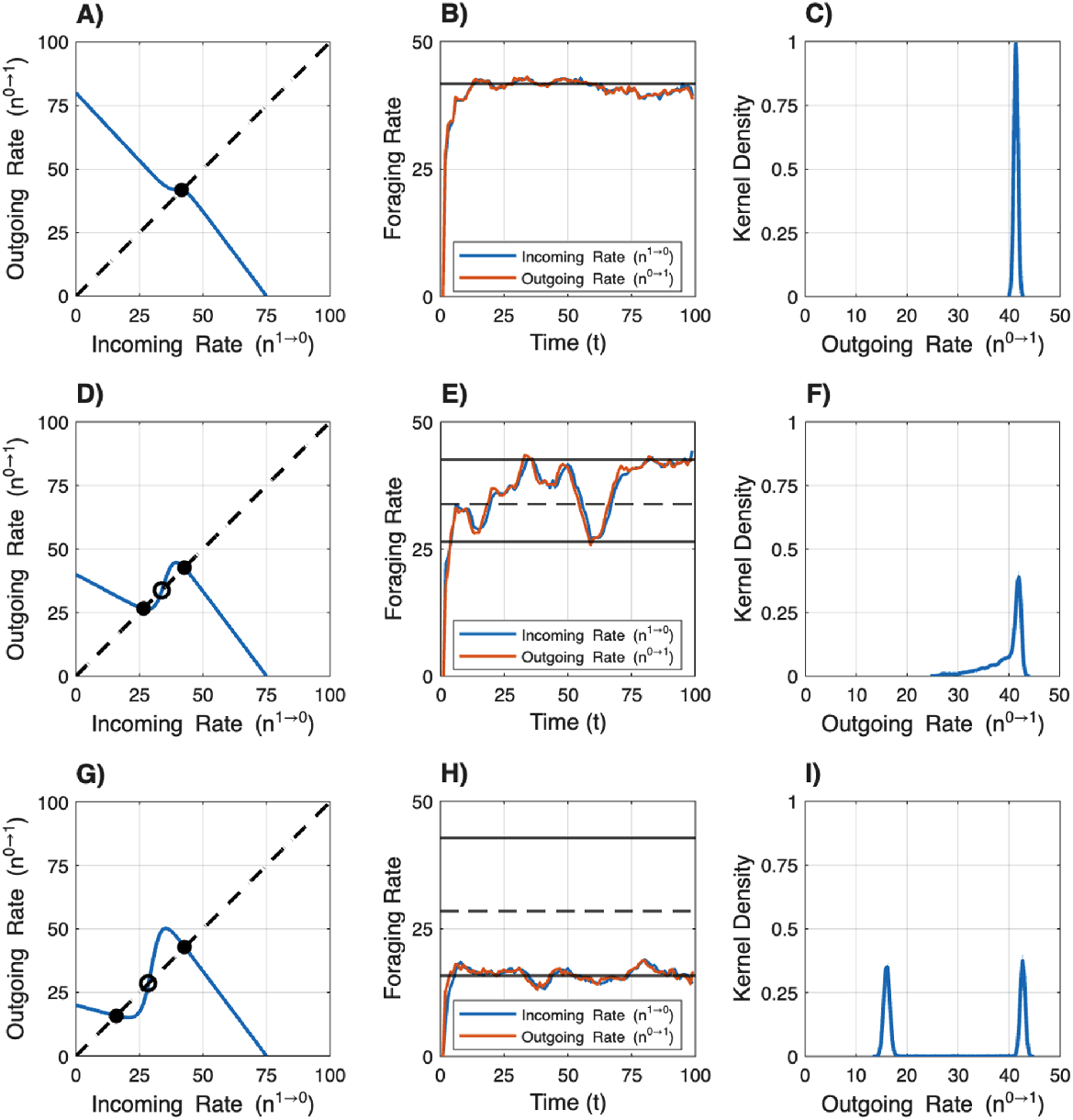
The flow-in model displays bistable dynamics. Rows correspond to simulations with different sets of parameters, and columns represent different types of measurements made for each simulation. Each of these graphs is derived from the identical flow-in model. The first column shows the numeric solutions for the final flow rate. The blue line shows all of the possible in and outflow rates for that combination of parameters, the dashed line gives the 1:1 relationship, and where these intersect gives the final flow rate of the system. Black points show stable steady states, and open circles give unstable steady states. In the second column, we have example simulations, showing how in and outflow rates change over time. Solid lines correspond to the steady states given as closed circles in the first column, while the dashed line represents the unstable steady state given in that column. The final column gives the kernel density of the final outgoing rate across many simulations, which is an approximation for the basin of attraction for the possible steady states.

The outflow rate for the density model is approximately linear with respect to the inflow rate, and so only 1 final flow rate is possible (Supporting Information S4).

### 5.2 Fisher Information

First, we measure the Fisher information with respect to *T*, which controls the sensitivity of the workers to the work signal for both the flow-in model and the density model. For high levels of sensitivity (low values of *T*), the resolution of the work signal is high and strongly influences the decision to forage. For low levels of sensitivity, environmental or internal noise disrupts the signal, and so the behavior of the workers is less determined by the strength of the signal.

Let 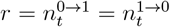, which is approximately true when the colony is at equilibrium and *p*(*r*|*T*) be the probability distribution of *r* given *T*. Let ℒ(*T*) be the Fisher information of the system with respect to *T*. Fisher information can be calculated as:

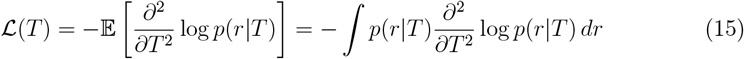

The second derivative of the log-likelihood can be expressed using the central difference formula [94]:

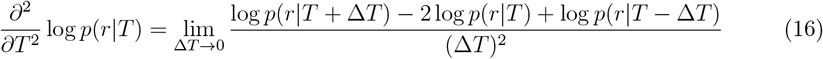

Substituting this into the definition of Fisher information yields:

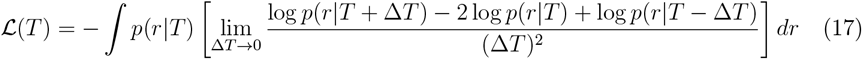

We estimate *p*(*r*|*T*), *p*(*r*|*T* + Δ*T*), and *p*(*r*|*T* − Δ*T*) at each combination of (*ϕ, µ*) by running simulations across all potential initial states 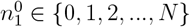 and then finding the average value of *r* in the latter half of the simulation.

We estimate the Fisher information with respect to *ρ*^1→0^, which sets the rate at which foraging ants return to the nest, using the same method.

### 5.3 Measuring Within and Across Simulation Variation and Probability of Attaining Different Steady States with Monte Carlo Simulations

At the onset of each simulation, 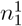 starts outside the nest and 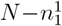 starts inside the nest. Each worker then uses the same transition probability matrix (equation 1) to transition into and out of the nest. All workers outside of the nest have the opportunity to go inside the nest first. The inflow rate of workers then determines the probability that workers inside the nest will flow outwards, so the probability that workers can leave the nest is only assessed after workers enter the nest. The simulation runs for *τ* timesteps. Every timestep, we count the number of workers that both leave and enter the nest. At the conclusion of each simulation, we measure the standard deviation of the outflow rate over *t*. For each combination of parameters, we run 100 simulations where the only difference is the number 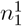 where 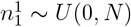. We therefore measure sensitivity to initial conditions as the standard deviation of the average outflow rate across these 100 replicates. Finally, we measure the probability of attaining different steady states when more than one solution is possible by counting the number of replications where the average simulated outflow rate is within 3 units of the expected higher steady state given the numeric approximation from section 2.3. All simulations were run in MATLAB [95].

### 5.4 Measuring Hysteresis

To measure hysteresis we systematically vary foraging potential *µ* over time so that its final value is equal to its initial value. To mimic the foraging behavior of single day [96], the foraging potential starts at a low value, indicating a period of low activity in the morning, the potential then peaks midday, and then is lowered again by the evening (Figure 5):

**Figure 5.**
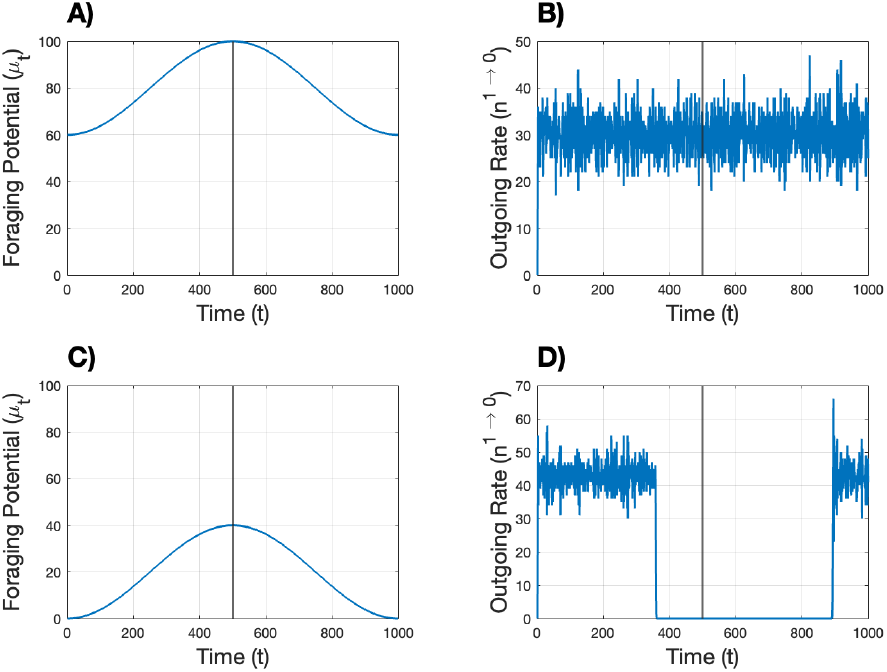
Flow-in model displays hysteresis for some parameter combinations. Simulations showing situations where there is hysteresis (A, B) and situations with no hysteresis (C, D). A and B show how *µ*_*t*_ changes over the course of the simulation, while B and D show how the outgoing rate of ants changes over time as a result of this change in *µ*_*t*_.

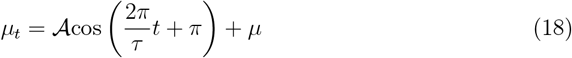

We then run simulations according to the same rules as those described in section For each timestep, we calculate the outflow of ants. After each simulation, we reflect the vector containing each outflow measurement across the point *τ/*2 and measure the cross-correlation between the first and second halves of the vector. A cross-correlation of 1 indicates perfect symmetry, and 0 indicates no symmetry. 1 minus the cross-correlation gives an estimate of the degree of hysteresis.

## Acknowledgments

The authors thank Dr. Deborah Gordon for her many helpful remarks on this manuscript. The authors also acknowledge Research Computing at Arizona State University for providing high-performance computing resources that have contributed to the research results reported within this paper.

## Supporting Information

### S1: Ising-Like Model of Foraging Behavior

A highly simplified model of foraging simply maps the current foraging rate *r*(*t*) to a future foraging rate *r*(*t*+*τ*) = *F*(*r*(*t*)). A continuous time version of this with a sigmoidal function *F* can be written as

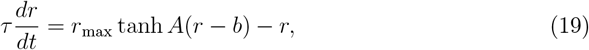

such that *r* approaches *F*(*r*) = *r*_max_ tanh *A*(*r* − *b*) over a timescale *τ*. Here, *A* defines the “amplification” of the signal when the “bias” *b* is zero.

This is reminiscent of the self-consistency equation of the mean-field Ising model. In particular, in an all-to-all coupled Ising model with Hamiltonian

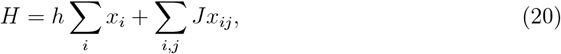

such that the probability distribution over states is given by 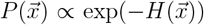 (here setting *β* = 1), the mean-field solution *x*_*i*_ = *m* ∀*i* satisfies

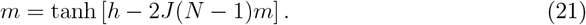

Therefore the equilibrium solution of (19) satisfies the same equation, with *r* = *m, A* = −2*J*(*N* − 1), and *b* = −*h/*(2*J*(*N* − 1)).

As a function of the amplification *A* and bias *b*, the Ising-like model has a cusp bifurcation, as can be seen by plotting a phase diagram (Figure S1, analogous to the third row of Figure 3).

**Figure S1.**
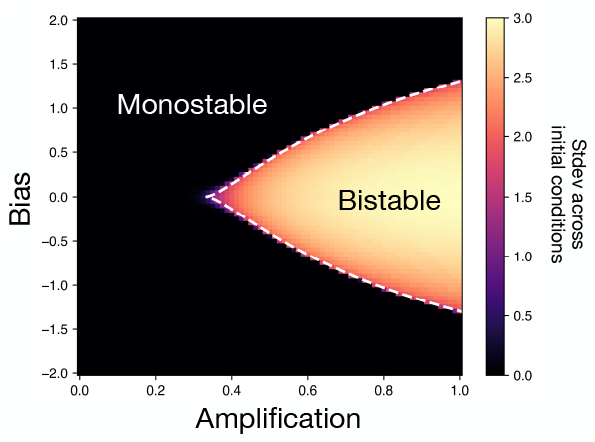
The Ising-like model has a cusp bifurcation. The standard deviation of final output rates in the Ising-like model, as a function of the effective amplification and bias parameters, shows a region of bistability surrounded by monostability. The edges of the bistable region correspond to saddle-node bifurcations (discontinuous transitions) that meet at a cusp bifurcation (continuous transition). These bifurcations can be traced numerically (exactly) in this model, as indicated by the dashed white lines.

### S2: Pagliara Model of Foraging Behavior

#### S2.1: Model Derivation

An existing detailed model of foraging dynamics in harvester ants includes both feedback from returning foragers and dynamics of recruitment within the nest [25]. We use this as an example of a model with more complicated dynamics that are more difficult to analyze mathematically.

In the detailed foraging model, returning ants add to a stimulus variable *s* that functions as a leaky integrator:

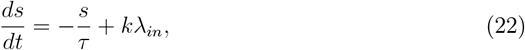

where *τ* is the leak timescale and *k* is the fixed amount by which each forager adds to the incoming signal (represented as delta functions in time in *λ*_*in*_). This signal then feeds into Fitzhugh–Nagumo (FN) oscillator dynamics, with each oscillation corresponding to one ant leaving the nest to forage. The oscillation dynamics are governed by

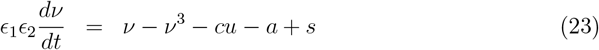

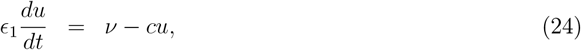

where *ϵ*_1_ and *ϵ*_2_ define the separation of timescales among *s, ν*, and *u*; *a* defines an offset to the stimulus *s*; and *c* defines a “volatility” via its control of both the propensity of the system to oscillate and the frequency of those oscillations. In particular, the FN system oscillates when the signal *s* has a value between two bifurcation points: *b*_1_ *< s < b*_2_, with 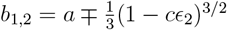.

We use mostly the default parameters as in Pagliara *et al*.: *k* = 0.3, *τ* = 0.41 s, *ϵ*_1_ = 0.2. We use a timescale separation for *ν* twice as large as in Pagliara *et al*., setting *ϵ*_2_ = 0.025, in order to simplify the model’s dependence on volatility *c*. We examine the model’s behavior with the original *ϵ*_2_ = 0.05 in the supplementary material.

In this parameter regime, increasing the volatility *c* increases the relative feedback strength due to an increase in the frequency of oscillations (and therefore in the rate of outgoing foragers for a given incoming rate). The *c* parameter also decreases the range of signals *s* that induce an oscillation, such that at very large *c*, no signals are able to induce oscillations.

In simulations, we start with initial rates of incoming ants set to one of {0.1, 0.25, 0.5, 1, 1.5, 2, 4, 6} ants/second for the first 11 minutes of the simulation, with incoming times chosen using a Poisson process. Outgoing ants return to the nest after a time chosen from a Poisson distribution with mean 10 minutes. We simulate 180 minutes of foraging dynamics using the MATLAB code provided in [25]. To generate input/output curves, we measure the rate of foraging given Poisson input with a constant mean rate that is then varied across simulations.

#### S2.2: Numerical Approximation of Steady State using Polynomials

Given the lack of a closed-form solution for fixed points in the [25] model, we cannot solve for the final flow rate directly. Additionally, the large computational loads of this model make it difficult to use the numeric approximations used in the Markovian model. Instead, for each combination of *c* and *a*, we fit a smooth curve to simulated outgoing foraging rates as a function of the incoming rate.

To identify an appropriate polynomial degree for modeling the relationship between incoming and outgoing ant traffic, we evaluated polynomial models of degree 1 through 8 using leave-one-out cross-validation (LOOCV; [97]). For each candidate degree, we computed the mean squared prediction error across all left-out observations and selected the optimal model using the 1-standard error (1-SE) rule [98].

Specifically, for each combination of *c* and *a*, we ran “open loop” simulations for a set of fixed incoming rates (0.1, 0.25, 0.5, 0.75, 1, 2, 3, 4, 5, 6 ants/s) and measured the resulting average outgoing rates over a period of *T* = 5 minutes. We then generated an orthogonal polynomial basis of increasing degree and computed the LOOCV error as follows: for each data point, a model was fit to the remaining *n* − 1 points, and the squared prediction error was calculated on the held-out point. The LOOCV error for a given degree was then defined as the mean of these squared errors. To avoid overfitting, we applied the 1-SE rule, which selects the most parsimonious model whose LOOCV error falls within one standard error of the minimum error observed. The standard error was computed from the LOOCV residuals at the degree with the lowest error. Among the degrees satisfying this criterion, the smallest was selected. This approach favors simpler models when they perform comparably to more complex alternatives.

Across all (*a, c*) combinations, the cubic polynomial (degree 3; Figure S2A) emerged as the most frequently selected model (Figure S2B) and exhibited strong average adjusted *R*^2^ performance (Figure S2C). Based on this systematic and conservative model selection procedure, we adopted the cubic polynomial as the canonical form for downstream analysis

**Figure S2.**
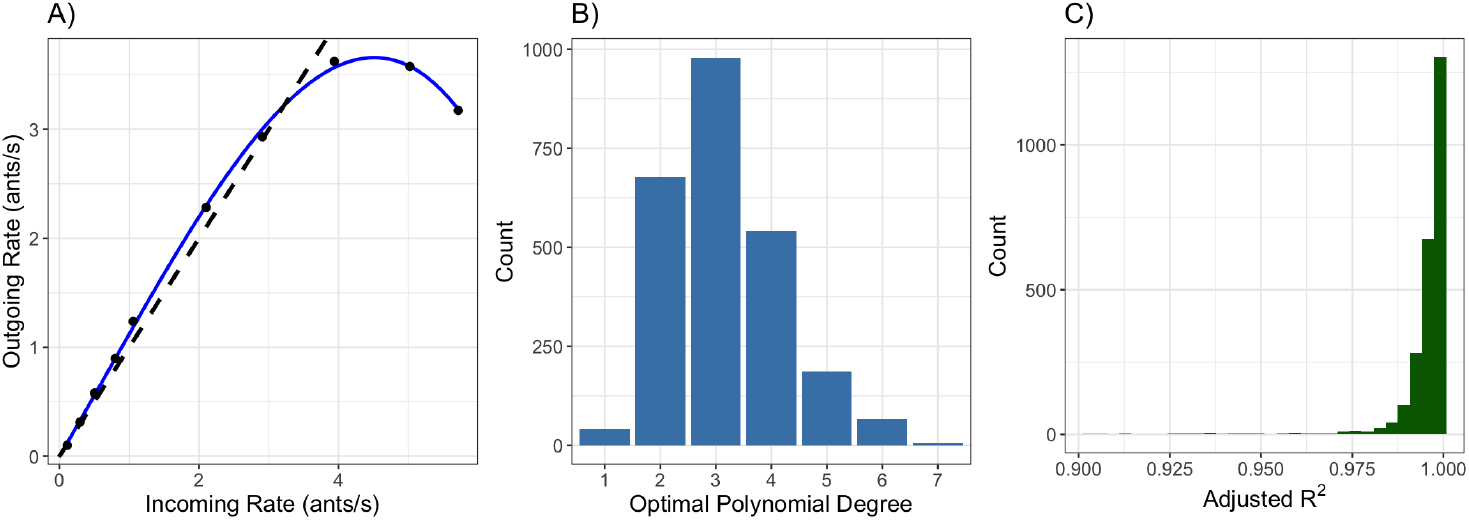
Fitting polynomials to self-consistency curves of the Pagliara model. A) Example cubic fit between incoming and outgoing foragers in the Pagliara model [25]. B) Distribution of optimal degree fits across all (*a, c*) combinations. C) Kernel density of adjusted *R*^2^ values for each dataset which has been fit using a cubic polynomial.

#### S2.3: Outlining Bistability Region in Parameter Space and Phase Diagram

Let *x* be the incoming rate of foragers in the Pagliara et al. model [25] and *f*(*x*) be the outgoing rate. We modeled the intersection between a cubic polynomial and the identity line using the expression

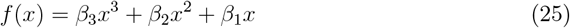

and examined the roots of the function

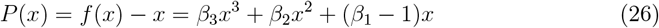

This cubic equation always has a root at *x* = 0, and we sought to identify regions in the (*a, c*) parameter space where *P*(*x*) additionally has a positive real double root, as this demarcates the edge of the bistable region.

To achieve this, we first fit third-order response surface models to the estimated polynomial coefficients *β*_1_, *β*_2_, and *β*_3_ as functions of the experimental parameters *a* and *c*.

We then evaluated the following algebraic conditions over a dense (*a, c*) grid to determine where *P*(*x*) exhibits a positive real double root:

1. The discriminant of the quadratic factor of *P*(*x*) is approximately zero:

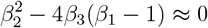

ensuring the existence of a repeated root.
2. The double root *r* is real and positive:

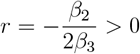

which requires that *β*_2_ and *β*_3_ have opposite signs.

Together, these conditions identify points in the (*a, c*) space where *P*(*x*) has exactly two real roots: a simple root at *x* = 0 and a repeated root at *x* = *r >* 0.

We evaluated the conditions numerically by computing the discriminant and root location across the fitted coefficient surfaces and plotting the contour where both criteria were satisfied, using a small numerical tolerance (*ε* = 10^−2^) to assess approximate equality.

In addition to identifying regions where the polynomial *P*(*x*) exhibits a positive real double root, we also examined where the original function *f*(*x*) has a slope at the origin greater than that of the identity line, and when *P*(*x*) has exactly one positive real root.

This corresponds to a situation when the polynomial *f*(*x*) crosses the origin with a slope larger than 1, such that it can only cross the identity line once for positive values, resulting in a single unstable steady state (at the origin) and a single steady state (where there is a positive real crossing). This gives us another boundary for the bistable region.

The slope of *f*(*x*) at the origin is

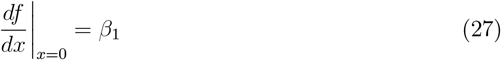

We evaluated the fitted response surface for *β*_1_(*a, c*) and identified regions in the (*a, c*) plane where

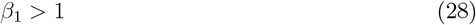

indicating that *f*(*x*) initially grows faster than the identity line *f*(*x*) = *x*. If we let *x*_1_ and *x*_2_ be the roots of *f*(*x*) − *x* (where *x*_0_ = 0), and we know that at least one of these roots are positive and the other negative, then by Vieta’s formula:

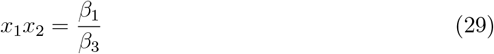

we know that the LHS must be negative, and since *β*_1_ *>* 1, *β*_3_ must be negative. Given these two conditions, we know that there must be at least one positive real root regardless of the value of *β*_2_. We know this because *f*(*x*) is positive for at least extremely small but positive values of *x* given the initial positive slope, and since lim_*x*→∞_ *f*(*x*) = −∞ when *β*_3_ *<* 0, by continuity there must be at least a single crossing across the identity line independent of the exact value of *β*_2_.

Analysis of stable states shows that this model, like the other versions of this foraging mechanism, displays a bifurcation in the phase space (Figure S3A). Again, like with the other models, this bistable region results in high variation in rates across initial conditions (Figure S3B). This region of bistability is well-bounded by the polynomial approximation of the self-consistency graph.

**Figure S3.**
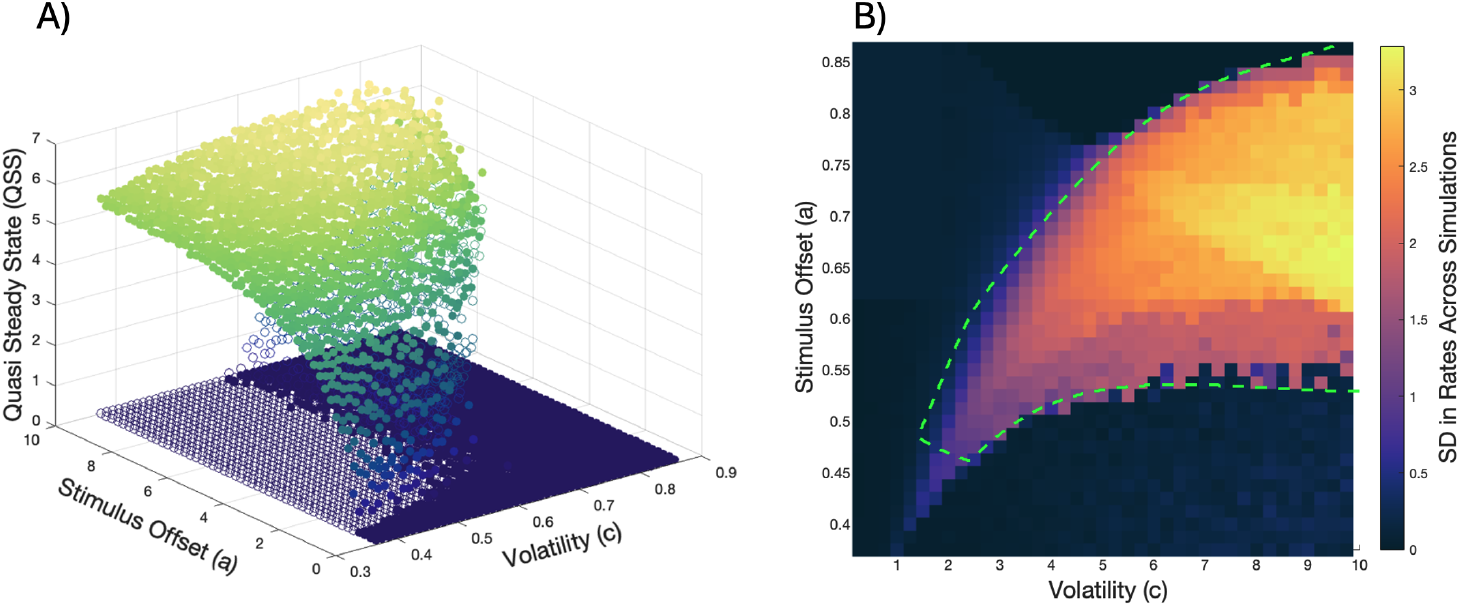
Bistability and bifurcations in the Pagliara model. A) Final flow rates (steady states) across combinations of values for the stimulus offset (*a*) and volatility (*c*). Closed circles show stable steady states, while open circles are unstable steady states. B) The standard deviation of outgoing/incoming rates of the Pagliara model across simulations with varying initial conditions. The green dashed line shows the region of bistability, as calculated in section B.3.

### S3: Measuring Effects of Parameters on the Flow-In Model

To ensure that the parameters in the composite function work as intended, we must show that *b* sets the intercept for equation 5, *µ* is the inflection point of the curve, and *T* still sets the steepness of the probability curve.

To find the intercept of equation 5, we set 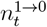 equal to 0:

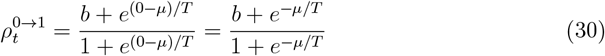

Clearly, the intercept is not exactly *ϕ*, but this function closely approximates *ϕ*. The slope of equation 6 with respect to *ϕ* is:

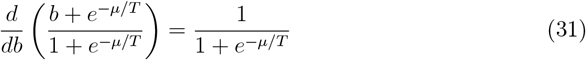

When *µ* ≫ *T, e*^−*µ/T*^ ≈ 0, so 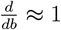. Conversely, when *µ* ≪ *T, e*^−*µ/T*^ ≈ 1, so 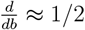. In this study, as we will vary *µ* across a wider range of parameters than *T*, as the range of *µ* will be set by the range of *N*, which will be constrained to the sizes of real *P. barbatus* colonies which have between 10 and 10,000 workers [99]. On the other hand, 0 *< T* ≤ 10, so for most simulations *µ* ≫ *T*, thus *ϕ* is extremely close to the intercept for most situations.

To find the inflection point of equation 5, we first find the second derivative of this equation with respect to 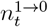:

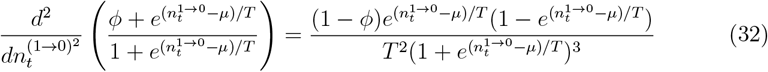

Setting this equation to 0 and then solving for 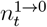 yields 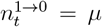, so *µ* is the inflection point.

To measure the effect of *T*, we can evaluate the slope of the function at the inflection point. The derivative of the unaltered version of equation 5 is the derivative of equation 3:

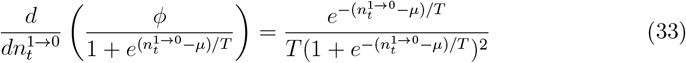

At the inflection point, 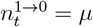, so if we substitute this in and simplify we find that the slope at the inflection point is 1*/*4*T*, so it only depends on T. Repeating this process for equation 5 gives a slope of (1 − *ϕ*)*/*4*T*, demonstrating that while *T* still determines the slope of this function, lower levels of foraging propensity will also influence it.

### S4: Monostability in Density Model

Bistability is not possible in the density model. The reason is that the presence of negative feedback (foragers leave the nest when there is a high density of workers, which subsequently decreases the probability that they leave) flattens the self-consistency curve, and thus there can only be a single steady state. We demonstrate this by showing that the inflow rate monotonically decreases with the number of ants within the nest while the outflow rate monotonically increases, therefore the two functions are only capable of crossing once.

Again, the probability that an ant leaves the nest at time *t* is given by:

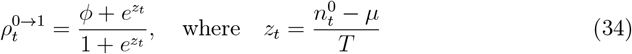

Thus, the expected outflow rate of ants from the nest at time *t* is:

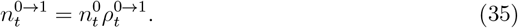

Conversely, ants outside the nest return with a fixed probability *ρ*^1→0^, yielding an inflow rate:

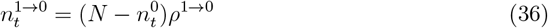

The derivatives of these rates with respect to 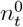 clarify their monotonicity. For the inflow rate, we have:

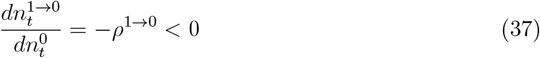

indicating a strictly decreasing relationship. For the outflow rate, the derivative is:

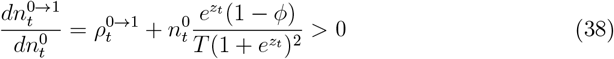

which is strictly positive for physically relevant parameter choices. Consequently, the self-consistency curve relating inflow to outflow can intersect the identity line 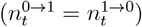 exactly once, ensuring a unique equilibrium solution for the foraging dynamics in this model.

### S5: Stability of Steady States in Flow-In Model at Various Steady States

We measure the stability of the model by calculating the absolute value of the derivative of the update function at each final flow rate where 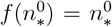. We calculate stability for all 3 steady states within the bistable region, and at the single steady state within the monostable region. At these final flow rates, the inflow and outflow rates are equal to their expected values:

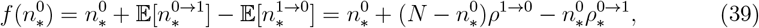

Where

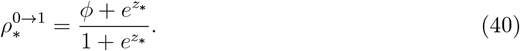

For the flow-in model,

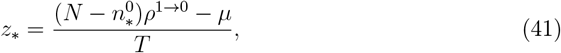

while for the density model,

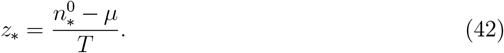

The derivative for the flow-in model is

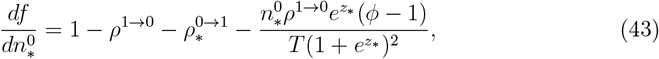

while for the density model,

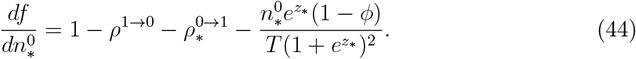

Finally, stability can be measured as 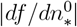 where 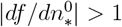 is unstable for this discrete-time map (Figure S5). We can see that the upper steady states are stable (Figure S5A) as well as all of the lower steady states (Figure S5C). However, each mid point is unstable (Figure S5B), meaning that within the bistable region, equilibrium can only be achieved for more than a few timesteps at the upper and lower steady states.

**Figure S5.**
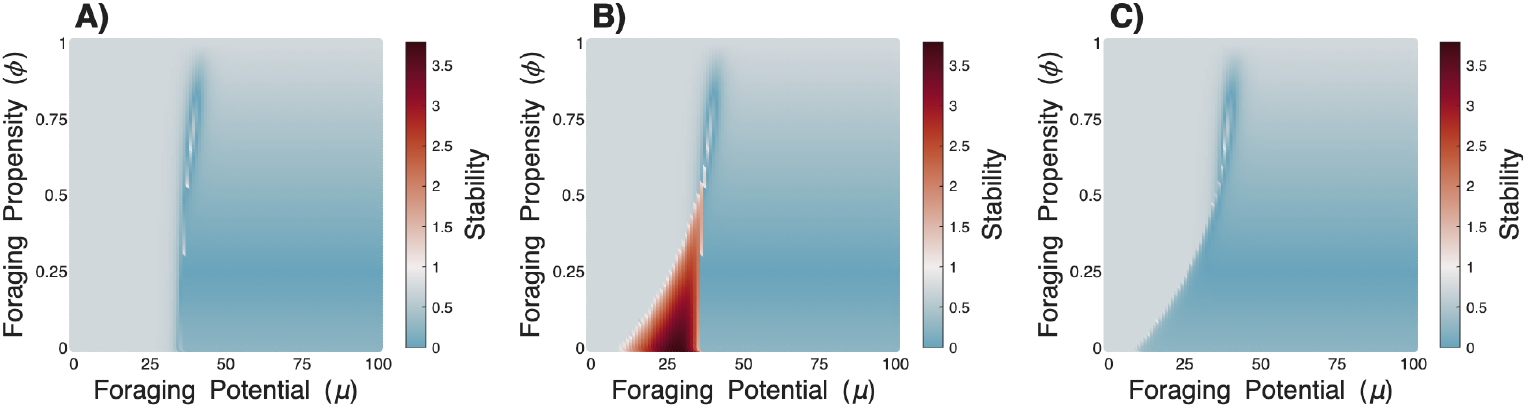
Upper and lower equilibria are stable, not intermediate equilibria in the identical flow-in model. Here we measure stability 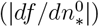 at each point in the phase space defined by *µ* and *ϕ*. This is measured independently at the upper (A), middle (B), and lower (C) equilibria for the identical flow-in model. Stability is never greater than 1 at the upper and lower steady states, but it is always greater than 0 at the intermediate state, making the latter unstable.

## Footnotes

1 This is closely related to the ambitions of catastrophe theory [26], which largely fell out of favor many decades ago after being over-generalized [27, 28] — we feel that similar ideas are worth revisiting in an era of big data, when we can more carefully test the limits of generalization

2 The renormalization group provides a powerful classification scheme of universality classes in the more specific case of continuous phase transitions [32, 33].

3 Recruitment within myrmecology is usually understood as “communication that brings nestmates to some point in *space* where work is required” [66, 67]. However, this term can have different definitions across different subfields of biology (i.e., in ecology it could describe the addition of early age classes to a population [68]). In the context of complex adaptive systems, we use a slightly more general definition, where recruitment is the process by which system units influence others to contribute to a particular *function* [69, 70].

4 Given the memoryless property of this model, this parameter also indirectly models the window of time workers within the nest will count incoming foragers to determine the rate.

5 However, biological constraints—such as communication bottlenecks (i.e. through changing the connectivity of neurons, [88]) —might preserve critical behavior by filtering environmental noise [89].

## References

1. Gordon DM. The ecology of collective behavior. PLoS biology 2014; 12:e1001805

2. Goldstone RL & Gureckis TM. Collective behavior. Topics in cognitive science 2009; 1:412–38

3. Tsuchiya M, Selvarajoo K, Piras V, Tomita M & Giuliani A. Local and global re-sponses in complex gene regulation networks. Physica A: Statistical Mechanics and its Applications 2009; 388:1738–46

4. Mojtahedi M, Skupin A, Zhou J, Castaño IG, Leong-Quong RYY, Chang H, Trachana K, Giuliani A & Huang S. Cell Fate Decision as High-Dimensional Critical State Transition. PLoS Biology 2016; 14:e2000640. doi: 10.1371/journal.pbio.2000640

5. Chen X, Randi F, Leifer AM & Bialek W. Searching for collective behavior in a small brain. Physical Review E 2019; 99:052418. doi: 10.1103/PhysRevE.99.052418

6. Daniels BC & Howard MW. Continuous Attractor Networks for Laplace Neural Manifolds. Computational Brain & Behavior 2025 :1–18

7. Cucker F & Smale S. Emergent behavior in flocks. IEEE Transactions on automatic control 2007; 52:852–62

8. Couzin ID. Collective cognition in animal groups. Trends in cognitive sciences 2009 Jan; 13:36–43. doi: 10.1016/j.tics.2008.10.002

9. Miller DL. Introduction to collective behavior and collective action. Waveland Press, 2013

10. Franks NR, Pratt SC, Mallon EB, Britton NF & Sumpter DJ. Information flow, opinion polling and collective intelligence in house–hunting social insects. Philosophical Transactions of the Royal Society of London. Series B: Biological Sciences 2002; 357:1567–83

11. Mizumoto N, Bardunias PM & Pratt SC. Parameter tuning facilitates the evolution of diverse tunneling patterns in termites. BioRxiv 2019 :836346

12. Schneider M, Bird AD, Gidon A, Triesch J, Jedlicka P & Cuntz H. Biological complexity facilitates tuning of the neuronal parameter space. PLoS computational biology 2023; 19:e1011212

13. Pratt SC & Sumpter DJ. A tunable algorithm for collective decision-making. Proceedings of the National Academy of Sciences 2006; 103:15906–10

14. Gordon DM. The ecology of collective behavior. en. Princeton Oxford: Princeton University Press, 2023

15. Solé R, Kempes CP, Corominas-Murtra B, De Domenico M, Kolchinsky A, Lachmann M, Libby E, Saavedra S, Smith E & Wolpert D. Fundamental constraints to the logic of living systems. en. Interface Focus 2024 Oct; 14:20240010. doi: 10.1098/rsfs.2024.0010

16. Mora T & Bialek W. Are biological systems poised at criticality? Journal of Statistical Physics 2011; 144:268–302

17. Shew WL, Yang H, Yu S, Roy R & Plenz D. Information capacity and transmission are maximized in balanced cortical networks with neuronal avalanches. Journal of neuroscience 2011; 31:55–63

18. Romanczuk P & Daniels BC. Phase transitions and criticality in the collective behavior of animals—self-organization and biological function. Order, Disorder and Criticality: Advanced Problems of Phase Transition Theory. World Scientific, 2023 :179–208

19. Muñoz MA. Colloquium: Criticality and dynamical scaling in living systems. Reviews of Modern Physics 2018; 90:031001

20. Attanasi A, Cavagna A, Del Castello L, Giardina I, Melillo S, Parisi L, Pohl O, Rossaro B, Shen E, Silvestri E et al. Finite-size scaling as a way to probe near-criticality in natural swarms. Physical review letters 2014; 113:238102

21. Hidalgo J, Grilli J, Suweis S, Munoz MA, Banavar JR & Maritan A. Information-based fitness and the emergence of criticality in living systems. Proceedings of the National Academy of Sciences 2014; 111:10095–100

22. Brush ER, Leonard NE & Levin SA. The content and availability of information affects the evolution of social-information gathering strategies. Theoretical Ecology 2016; 9:455–76

23. Poel W, Daniels BC, Sosna MMG, Twomey CR, Leblanc SP, Couzin ID & Ro-manczuk P. Subcritical escape waves in schooling fish. Science Advances 2022; 8. Publication Title: Sci. Adv Volume: 8:eabm6385

24. Sarkanych P, Krasnytska M, Gómez-Nava L, Romanczuk P & Holovatch Y. Individual bias and fluctuations in collective decision making: from algorithms to Hamiltonians. Physical Biology 2023; 20:045005

25. Pagliara R, Gordon DM & Leonard NE. Regulation of harvester ant foraging as a closed-loop excitable system. PLoS computational biology 2018; 14:e1006200

26. Thom R. Structural Stability, Catastrophe Theory, and Applied Mathematics. en. SIAM Review 1977 Apr; 19:189–201. doi: 10.1137/1019036

27. Kolata GB. Catastrophe Theory: The Emperor Has No Clothes. en. Science 1977

28. Zahler RS & Sussmann HJ. Claims and accomplishments of applied catastrophe theory. en. Nature 1977 Oct; 269:759–63. doi: 10.1038/269759a0

29. Daniels BC, Laubichler MD & Flack JC. Introduction to the special issue: quantifying collectivity. 2021

30. Crawford JD. Introduction to bifurcation theory. Reviews of modern physics 1991; 63:991

31. Strogatz SH. Nonlinear Dynamics and Chaos: With Applications to Physics, Biology, Chemistry, and Engineering. CRC Press, 2015

32. Goldenfeld N. Lectures on Phase Transitions and the Renormalization Group. Westview Press, 1992

33. Sethna J. Entropy, order parameters, and complexity. Oxford University Press, 2006

34. Rand DA, Raju A, Sáez M, Corson F & Siggia ED. Geometry of gene regulatory dynamics. en. Proceedings of the National Academy of Sciences 2021 Sep; 118:e2109729118. doi: 10.1073/pnas.2109729118

35. Krakauer DC & Flack JC. Better living through physics. Nature 2010; 467:661–1

36. Vicsek T, Czirók A, Ben-Jacob E, Cohen I & Shochet O. Novel type of phase transition in a system of self-driven particles. Physical review letters 1995; 75:1226

37. Baglietto G & Albano EV. Nature of the order-disorder transition in the Vicsek model for the collective motion of self-propelled particles. Physical Review E—Statistic Nonlinear, and Soft Matter Physics 2009; 80:050103

38. Wang SH, Siebenhühner F, Arnulfo G, Myrov V, Nobili L, Breakspear M, Palva S & Palva JM. Critical-like brain dynamics in a continuum from second-to first-order phase transition. Journal of Neuroscience 2023; 43:7642–56

39. Mukherjee G & Manna S. Phase transition in a directed traffic flow network. Physical Review E—Statistical, Nonlinear, and Soft Matter Physics 2005; 71:066108

40. Szabo B, Szöllösi G, Gönci B, Jurányi Z, Selmeczi D & Vicsek T. Phase transition in the collective migration of tissue cells: experiment and model. Physical Review E—Statistical, Nonlinear, and Soft Matter Physics 2006; 74:061908

41. Mitesser O, Weissel N, Strohm E & Poethke HJ. Adaptive dynamic resource allocation in annual eusocial insects: environmental variation will not necessarily promote graded control. BMC ecology 2007; 7:1–13

42. Daniels BC, Ellison CJ, Krakauer DC & Flack JC. Quantifying collectivity. Current opinion in neurobiology 2016; 37:106–13

43. Noori HR & Noori HR. Examples of Hysteresis Phenomena in Biology. Springer, 2014

44. Guo X, Lin MR, Azizi A, Saldyt LP, Kang Y, Pavlic TP & Fewell JH. Decoding alarm signal propagation of seed-harvester ants using automated movement tracking and supervised machine learning. Proceedings of the Royal Society B 2022; 289:20212176

45. Lin MR, Guo X, Azizi A, Fewell JH & Milner F. Mechanistic modeling of alarm signaling in seed-harvester ants. Mathematical Biosciences and Engineering 2024; 21:5536–55

46. Beekman M, Sumpter DJ & Ratnieks FL. Phase transition between disordered and ordered foraging in Pharaoh’s ants. Proceedings of the National Academy of Sciences 2001; 98:9703–6

47. Hein AM, Rosenthal SB, Hagstrom GI, Berdahl A, Torney CJ & Couzin ID. The evolution of distributed sensing and collective computation in animal populations. Elife 2015; 4:e10955

48. Sumpter DJ & Pratt SC. Quorum responses and consensus decision making. Philosophical transactions of the Royal Society B: biological sciences 2009; 364:743–53

49. Wilson EO. Chemical communication among workers of the fire ant Solenopsis saevissima (Fr. Smith) 1. The organization of mass-foraging. Animal behaviour 1962; 10:134–47

50. Baudier K & O’Donnell S. Structure and thermal biology of subterranean army ant bivouacs in tropical montane forests. Insectes Sociaux 2016; 63:467–76

51. Hölldobler B & Wilson EO. The ants. Harvard University Press, 1990

52. Kay A. Applying optimal foraging theory to assess nutrient availability ratios for ants. Ecology 2002; 83:1935–44

53. Greene MJ & Gordon DM. Interaction rate informs harvester ant task decisions. Behavioral Ecology 2007; 18:451–5

54. Gordon DM. The rewards of restraint in the collective regulation of foraging by harvester ant colonies. Nature 2013; 498:91–3. doi: 10.1038/nature12137

55. Pinter-Wollman N, Bala A, Merrell A, Queirolo J, Stumpe MC, Holmes S & Gordon DM. Harvester ants use interactions to regulate forager activation and availability. Animal behaviour 2013; 86:197–207

56. Davidson JD, Arauco-Aliaga RP, Crow S, Gordon DM & Goldman MS. Effect of interactions between harvester ants on forager decisions. Frontiers in ecology and evolution 2016; 4:115

57. Udiani O, Pinter-Wollman N & Kang Y. Identifying robustness in the regulation of collective foraging of ant colonies using an interaction-based model with backward bifurcation. Journal of theoretical biology 2015; 367:61–75

58. Feng T & Kang Y. Foraging dynamics in social insect colonies: Mechanisms of backward bifurcations and impacts of stochasticity. Mathematical Biosciences 2025 :109436

59. Traniello JF. Foraging strategies of ants. Annual review of entomology 1989; 34:191– 210

60. Cao TT. High social density increases foraging and scouting rates and induces polydomy in Temnothorax ants. Behavioral Ecology and Sociobiology 2013; 67:1799– 807

61. Somogyi AÁ, Tartally A, Maák IE & Barta Z. Colony size, nestmate density and social history shape behavioural variation in Formica fusca colonies. Ethology 2020; 126:727–34

62. Dawson DA. Critical dynamics and fluctuations for a mean-field model of cooperative behavior. Journal of Statistical Physics 1983; 31:29–85

63. Wang W, Chen Y & Huang J. Heterogeneous preferences, decision-making capacity, and phase transitions in a complex adaptive system. Proceedings of the National Academy of Sciences 2009; 106:8423–8

64. Garnier S, Gautrais J & Theraulaz G. The biological principles of swarm intelligence. Swarm intelligence 2007; 1:3–31

65. Cohen IR. Tending Adam’s Garden: evolving the cognitive immune self. Elsevier, 2000

66. Wilson EO. The insect societies. 1971

67. Planqué R, Van Den Berg JB & Franks NR. Recruitment strategies and colony size in ants. PLoS One 2010; 5:e11664

68. Nishizaki MT & Ackerman JD. Settlement and recruitment of pelagic larvae to benthic habitats. Encyclopedia of water: Science, technology, and society 2019 :1– 15

69. Muñoz-Nava LM, Flores-Flores M & Nahmad M. Inducing your neighbors to become like you: cell recruitment in developmental patterning and growth. The International Journal of Developmental Biology 2021; 65:357–64

70. Nozari E & Cortés J. Selective recruitment in hierarchical complex dynamical networks with linear-threshold rate dynamics. 2018 IEEE Conference on Decision and Control (CDC). IEEE. 2018 :5227–32

71. Fajgenbaum DC & June CH. Cytokine storm. New England Journal of Medicine 2020; 383:2255–73

72. Gordon DM. The ecology of collective behavior. 2023

73. Prokopenko M, Lizier JT, Obst O & Wang XR. Relating Fisher information to order parameters. Physical Review E 2011 Oct; 84:041116. doi: 10.1103/PhysRevE.84.041116

74. Loengarov A & Tereshko V. Phase transitions and bistability in honeybee foraging dynamics. 2008

75. Klamser PP & Romanczuk P. Collective predator evasion: Putting the criticality hypothesis to the test. PLoS computational biology 2021; 17:e1008832

76. Scarpetta S, Apicella I, Minati L & De Candia A. Hysteresis, neural avalanches, and critical behavior near a first-order transition of a spiking neural network. Physical Review E 2018; 97:062305

77. Cinquin O & Demongeot J. Roles of positive and negative feedback in biological systems. Comptes rendus. Biologies 2002; 325:1085–95

78. Feinerman O & Korman A. Individual versus collective cognition in social insects. Journal of Experimental Biology 2017; 220:73–82

79. Buffin A, Sasaki T & Pratt SC. Scaling of speed with group size in cooperative transport by the ant Novomessor cockerelli. PloS one 2018; 13:e0205400

80. Schatz B, Lachaud JP & Beugnon G. Graded recruitment and hunting strategies linked to prey weight and size in the ponerine ant Ectatomma ruidum. Behavioral Ecology and Sociobiology 1997; 40:337–49

81. McCreery HF, Gemayel G, Pais AI, Garnier S & Nagpal R. Hysteresis stabilizes dynamic control of self-assembled army ant constructions. Nature communications 2022; 13:1160

82. Hidalgo J, Grilli J, Suweis S, Maritan A & Munoz MA. Cooperation, competition and the emergence of criticality in communities of adaptive systems. Journal of Statistical Mechanics: Theory and Experiment 2016; 2016:033203

83. Bagley RJ, Farmer JD, Kauffman SA, Packard NH, Perelson AS & Stadnyk I. Modeling adaptive biological systems. Biosystems 1989; 23:113–37

84. Cassidy AS, Merolla P, Arthur JV, Esser SK, Jackson B, Alvarez-Icaza R, Datta P, Sawada J, Wong TM, Feldman V et al. Cognitive computing building block: A versatile and efficient digital neuron model for neurosynaptic cores. The 2013 international joint conference on neural networks (IJCNN). IEEE. 2013 :1–10

85. Stožer A, Marković R, Dolenšek J, Perc M, Marhl M, Slak Rupnik M & Gosak M. Heterogeneity and delayed activation as hallmarks of self-organization and criticality in excitable tissue. Frontiers in Physiology 2019; 10:869

86. Moretti P & Muñoz MA. Griffiths phases and the stretching of criticality in brain networks. Nature communications 2013; 4:2521

87. Bray A. Nature of the Griffiths phase. Physical review letters 1987; 59:586

88. Kinouchi O & Copelli M. Optimal dynamical range of excitable networks at criticality. Nature physics 2006; 2:348–51

89. Zhu K, Feng X, Du X, Gu Y, Yu W, Wang H, Chen Q, Chu Z, Chen J & Qin B. An information bottleneck perspective for effective noise filtering on retrieval-augmented generation. arXiv preprint 2406.01549 2024

90. Feng T, Liu C & Milne R. Noise-Driven Transitions in Collective Foraging of Ant Colonies. Bulletin of Mathematical Biology 2025; 87:1–46

91. Kuznetsov YA. Elements of Applied Bifurcation Theory. New York: Springer-Verlag, 1995

92. Gordon DM, Dektar KN & Pinter-Wollman N. Harvester ant colony variation in foraging activity and response to humidity. PloS one 2013; 8:e63363

93. Friedman DA, Pilko A, Skowronska-Krawczyk D, Krasinska K, Parker JW, Hirsh J & Gordon DM. The role of dopamine in the collective regulation of foraging in harvester ants. Iscience 2018; 8:283–94

94. Arehart E, Jin T & Daniels BC. Locating decision-making circuits in a heterogeneous neural network. Frontiers in Applied Mathematics and Statistics 2018; 4:11

95. Inc. TM. MATLAB version: 9.14.0 (R2023a). Natick, Massachusetts, United States, 2023. Available from: https://www.mathworks.com

96. Gordon DM. The dynamics of the daily round of the harvester ant colony (Pogonomyrmex barbatus). Animal Behaviour 1986; 34:1402–19

97. Wong TT. Performance evaluation of classification algorithms by k-fold and leaveone-out cross validation. Pattern recognition 2015; 48:2839–46

98. Bulteel K, Mestdagh M, Tuerlinckx F & Ceulemans E. VAR (1) based models do not always outpredict AR (1) models in typical psychological applications. Psychological methods 2018; 23:740

99. Gordon DM. How colony growth affects forager intrusion between neighboring harvester ant colonies. Behavioral ecology and sociobiology 1992; 31:417–27

